# Lipid nanoparticle induced model endosomal membrane disruption

**DOI:** 10.64898/2026.09.11.751062

**Authors:** Shreya Pramanik, Aishwarya Vasudevan, Zhenghua Liang, Archana Karole, Mathieu Berchel, Paul-Alain Jaffrès, Yulia Eygeris, Gaurav Sahay

## Abstract

Endosomal escape remains the central bottleneck limiting lipid nanoparticle (LNP)-mediated mRNA delivery. In this study, we use giant unilamellar vesicles (GUVs) mimicking the endosomal membrane to directly visualize the interaction and membrane disruptions caused by two model LNPs constituted by two different ionizable lipids: LP01 and BiP-20 (a Branched ionizable Phospholipid) that are both benchmark ionizable lipids for in vivo gene editing in liver. Despite identical TNS-measured apparent pKa (6.6) for both lipids in our conditions, LP01 LNP showed maximal membrane engagement at pH 5.8, while BiP20 LNP required pH 5.1, demonstrating that bulk pKa alone cannot predict the pH at which functional membrane interaction occurs. Using confocal microscopy, we show that the LNP engagement with the endosomal membrane proceeds through first electrostatic attraction, clustering, membrane disruption by forming highly curved morphologies, and finally leading to the rupture of the whole endosomal mimicking membrane. Notably, the mRNA cargo is released only upon complete lysis of the membrane. This indicates that the membrane destabilization and cargo release are mechanistically distinct steps. These findings establish a cell-free platform for dissecting individual sub-steps of endosomal escape.

## INTRODUCTION

Intracellular delivery of nucleic acids is a cornerstone of modern therapeutic development, particularly in the context of mRNA-based vaccines, gene therapy and cancer therapeutics. Lipid nanoparticles (LNPs) have emerged as the leading non-viral delivery platform for this purpose, demonstrating high clinical efficacy and their deployment in FDA approved mRNA vaccines^1^. LNPs are undergoing clinical trials for gene therapy and cancer, with promising results^2^. The LNPs are engulfed by the cells by endocytosis and get trapped in a membrane-bound cell organelle called an endosome. Endosomes are characterized by the anionic lipid composition of their membrane, and their progressively acidic pH ranging from approximately 6.0–6.5 in early endosomes to 4.5–5.0 in late endosomes and lysosomes. For therapeutic efficacy, the nucleic acid cargo of the LNPs should escape the endosomes before the endosomes mature into lysosomes. This process of endosomal escape is the bottleneck in LNP-mediated intracellular nucleic acid delivery^3,4^. Studying the molecular and biophysical mechanism of endosomal escape will help in the rationale behind the designs of next-generation delivery systems with improved therapeutic efficacy^5^.

LNP shell is made of four major components—ionizable lipids, phospholipids, cholesterol, and PEG lipids—that encapsulate nucleic acid cargo. The extracellular environment and cytosol are maintained at a physiological pH of approximately 7.4, at which ionizable lipids remain mostly unprotonated and lipid nanoparticles (LNPs) are effectively uncharged. Upon cellular uptake, LNPs become trapped within endosomes, which are characterized by a progressively acidifying lumen ^6,7^. This acidification triggers protonation of the ionizable lipid component, generating a net positive charge on the nanoparticle surface. Charge acquisition is considered a critical step for LNP engagement with the endosomal membrane. The endosomal membrane is enriched in negatively charged phospholipids relative to the plasma membrane^8^, and this anionic character electrostatically attracts the now-cationic LNP, promoting membrane association and subsequent destabilization^9,10^. If this destabilization results in cargo release into the cytosol prior to lysosomal degradation, the RNA payload can be translated into protein.

Studying LNP-membrane interactions in the native intracellular environment has experimental and imaging challenges^11,12^. The endosomal compartments are dynamic and compositionally complex, making it difficult to isolate and characterize individual molecular events at sufficient spatial and temporal resolution. There are studies using lipid monolayers^13^, supported lipid bilayers^14,15^ and computational simulations^16,17^ to study the mechanism of endosomal escape, but these have limitations related to membrane curvature or short time scales. In this study, we use Giant Unilamellar Vesicles (GUVs), a biomimetic lipid membrane system, to mimic the endosomal lipid composition. GUVs are made of a lipid bilayer that closes upon itself and separates the aqueous media into two regions, inside and the outside of the GUV. Their diameter is typically in micrometers and can be easily observed using light microscopy. This offers an advantage over the nanometer sized endosomes in the cells, which require substantial technical efforts to be isolated or imaged^18,19^.

Early endosomes are enriched in phosphatidylcholine (PC), phosphatidylethanolamine (PE), phosphatidylserine (PS), phosphatidylinositol 3-phosphate (PI3P), and cholesterol^8,20,21^. All these components have different roles in maintaining the functionality of the endosome. The PC and PE are lipids with neutral head groups, while the PS and PI3P have negatively charged head groups, thus making the early endosomal membrane anionic. PI3P is generated on the early endosomal membrane by the lipid kinase VPS34, and it serves as a critical signaling lipid that recruits proteins involved in endosomal sorting and trafficking^22^. Cholesterol plays a critical role in regulating membrane fluidity, rigidity, and the formation of lipid microdomains. We used a similar lipid composition to make the GUVs to mimic the early endosomal membranes and to study their interactions with the LNPs.

Here, we employed confocal fluorescence microscopy with a GUV model membrane system to investigate the interaction between LNPs and lipid bilayers to provide a mechanistic insight into endosomal escape. We first optimized the lipid composition of a model endosomal membrane (POPC:DOPE:DOPS:Chol:PI3P, 55.25:20.25:10.25:10.25:4 mol%) and established an image analysis protocol to quantify the interaction between LNP and GUV. We captured a discrete morphological sequence starting with LNP clustering on the membrane, RNA exposure through hemifusion-like intermediates, and eventual GUV lysis. This traces how the incorporation of ionizable lipid in the lipid membrane drives endosomal membrane destabilization and ultimately triggers mRNA release into the cytoplasm.

## RESULTS AND DISUSSSION

### Quantitative analysis of membrane intensity and optimization of membrane composition of endosomal model membrane

GUVs are spherical, micrometer-sized closed lipid membranes enclosing aqueous solution, and keeping their spherical shape requires matching the osmolarity of the interior and exterior solutions. In our experiments, the GUVs are prepared in 2mM HEPES and 300mM sucrose. This concentration was chosen to match the osmolarity of PBS in which the LNPs are suspended. To establish a lower pH on the outside, GUV containing solution is mixed with PBS at a lower pH. The final pH of the GUV solution was measured using a pH meter equipped with a microprobe. PBS has low density compared to sucrose, thus making the GUVs denser. This helps settle the GUVs on the coverslip, making it possible to image them using an inverted microscope.

To establish an endosomal model membrane for our experiments, we optimized the lipid composition of the GUVs. Selecting the optimal composition required ensuring the biological relevance of each lipid component without introducing too many variables that could confound the results. We started with the simplest bilayer composition, POPC:Cholesterol (80:20 mol%), and exposed it to LNPs doped with Rh-DHPE dye at pH 5.8. The LNPs showed a very low signal at the membrane. POPC forms the backbone of the lipid bilayer, and 20 mol% cholesterol represents the high cholesterol content of the endosomal membrane; both components are uncharged, making the bilayer neutral, so the positively charged LNPs at low pH did not interact much with it.

Introducing 10 mol% of the negatively charged phospholipid POPS increased the electrostatic interaction between the membrane and the protonated LNPs. Finally, we added DOPE and PI(3)P to more closely resemble the native early endosomal lipid composition. The composition POPC:Chol:DOPS:DOPE:PI3P at 55.25:10.25:10.25:20.25:4 mol% (identified below as endosomal composition) showed the maximum interaction with the LNPs, and we used it for all further experiments. Our decision to include and focus on the PI3P lipid was motivated by in vivo LysoTag immunoprecipitation experiments, in which proteomic profiling of the isolated endolysosomal compartment after LNP treatment revealed a pronounced remodeling of the organelle, including depletion of the early endosomal marker EEA1 and its downstream trafficking machinery. Because EEA1 is recruited to the endosomal membrane through FYVE-domain recognition of PI3P, this loss pointed to a perturbation of the PI3P pool itself and led us to hypothesize that ionizable lipids directly engage PI3P-enriched domains, motivating the fundamental lipid-lipid interaction studies in GUVs described here^23^.

We have selected two LNPs that include either the ionizable lipid LP01 (composition: LP01/Cholesterol/DSPC/DMG-PEG2000: 45.5/44/9/1.5 mol%) or the branched ionizable phospholipid BiP-20 (BiP20/Cholesterol/DOPE/DMPE-PEG2000: 50/38.5/9.5/1.5 mol%) These LNPs correspond to optimized formulations reported previously (LP01^24^, BiP-20^23^) and that are used for in vivo gene editing in the liver. LP01 is in phase 3 clinical trials (NCT06128629, NCT06672237) and recent results demonstrate that BiP-20 outperformed LP01 for liver gene editing in a murine model [50]. When these LNPs are suspended in an acidic buffer, the ionizable lipids become mostly protonated and interact with the negatively charged GUV membranes. Instead of a uniform distribution across the GUV membrane, we observe local clustering of the LNPs, on the GUV surface. However, there was a uniform fluorescence signal of Rh-DHPE throughout the GUV membrane, showing uniform mixing of the helper lipid from the LNPs. This clustering of LNPs is consistent with local membrane defects recruiting other particles to the same position^25–28^. Initial LNP adhesion and lipid insertion at a given site generate localized packing defects and curvature stress. Such packing defects expose the hydrophobic part of the lipid bilayer and increase the recruitment of the LNPs at the same area, propagating the formation of LNP-GUV clusters. Once in the membrane, the helper lipids mix with the endosomal membrane while the ionizable lipids remain at the point of contact. This has also been shown by Kjølbye et al in the molecular dynamics simulations^16^.

As shown in Figure 1, we quantified the fluorescence of the membrane. To avoid fluorescence signal artifacts, we selected portions of the GUV without any visible puncta. It is to be noted that in some cases, the puncta were not visible in the image of the GUV. This is because the LNPs interact with the GUV at different planes, while the images are taken at the plane where the middle of GUVs. The fluorescence of fluorophores can be affected by surrounding pH^29–31^ and that could cause other artifacts in the experiments. To avoid that, we take the ratio of the fluorescence of the membrane to that of the background.

**Figure 1:**
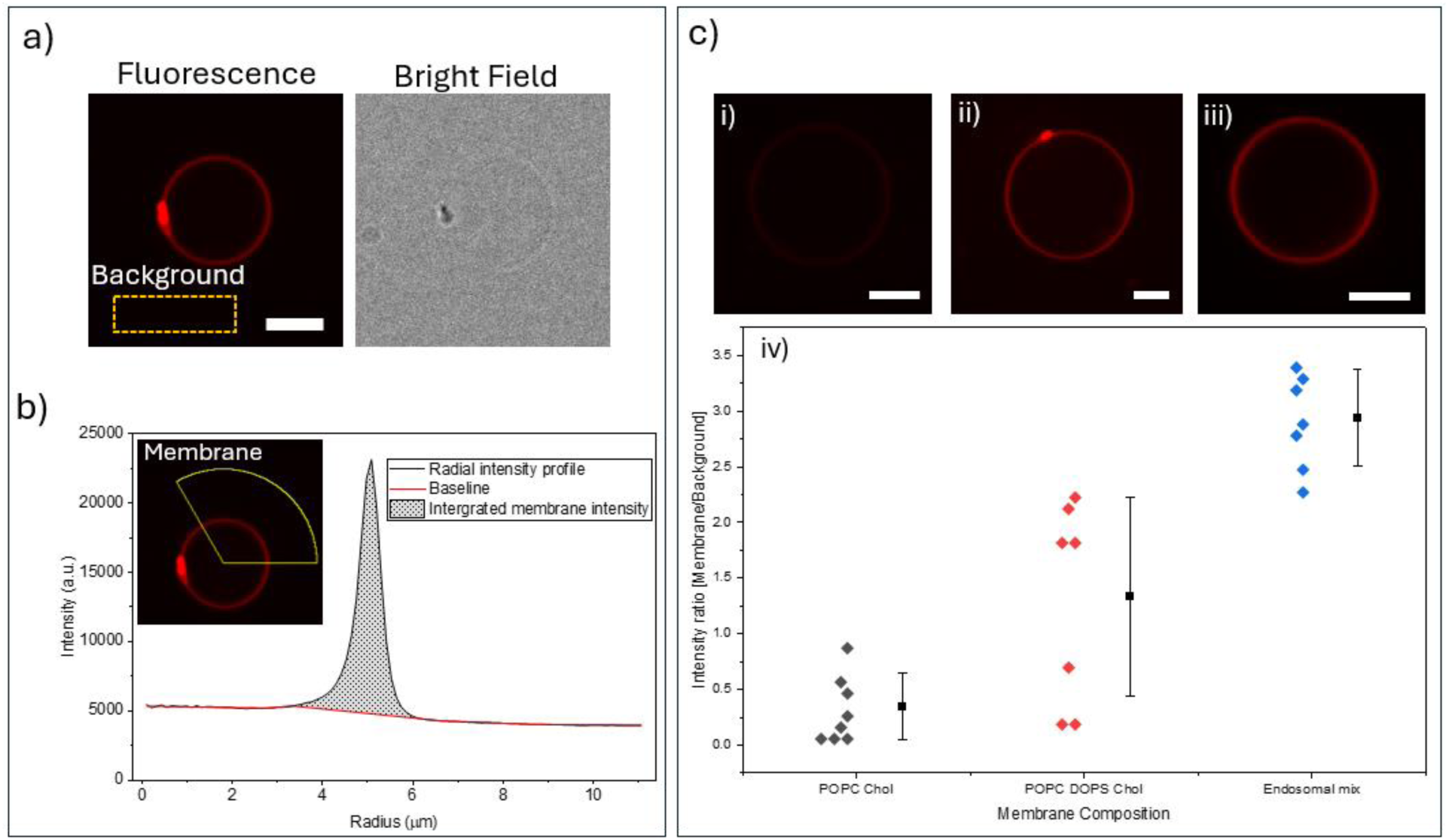
GUVs with different lipid compositions, POPC: Chol (80:20 mol%), POPC: Chol: POPS (70:20:10 mol%) and Endosomal mix, were exposed to LP01 LNPs doped with fluorescent lipid Rh-DHPE, encapsulating fluc mRNA, with the final mRNA concentration in all samples is 720ng at pH 5.8. a) Selecting a region of interest (ROI) around the GUV avoiding lipid clusters. n=3 such ROIs were selected, and their average intensity was set as the ‘Background Intensity’. b) Part of the GUV membrane was selected avoiding membrane clusters, its fluorescence intensity across the radius was plotted (black) and the baseline (Origin) was subtracted. The integrated area of the subtracted region (shaded region) was set as the ‘Membrane Intensity’. c) The ratio of the intensities (Membrane/ Background) was calculated for each composition. i) to iii) show representative images of the three compositions. iv) Quantitative intensity ratios measured for the interaction of various lipid compositions on interacting with LP01 LNPs. Each of the diamond points corresponds to data from 1 GUV. Each solid black square and the whiskers correspond to mean and standard deviation of GUVs. All scale bars: 5 µm

### Lipid Nanoparticles with similar TNS pKa interact with the membrane differently

The apparent pKa of the LNP is generally measured with the TNS fluorescence assay, a widely used assay to correlate lipid ionization properties with nanoparticle potency. However, a recent study by McMillan et al has shown that although pKa is an important parameter influencing the ionization behavior of the LNP, it cannot predict the performance and functionality of the LNP *in vitro* or *in vivo*^32^. This is likely because the TNS assay measures the apparent surface properties of LNPs but fails to capture the intricacies of core lipid protonation and the molecular behavior of the LNP when interacting with the endosomal membrane^33^. For the same ionizable lipids, the apparent pKa can be different based on factors such as the type of helper lipids^34–36^, tail length and symmetry of the ionizable lipids^37–41^, packaging^42^, and RNA cargo^43^. Not just the pKa, but the tail lengths of ionizable lipids also play a major role in delivery of their cargo^38,44^. The tail lengths and structures of the ionizable lipids govern how easily they adapt into highly curved surfaces such as hexagonal phases^45^. To investigate how these factors may affect the molecular interactions with model endosomal membranes, we prepared and compared the behavior of two LNP populations with different ionizable lipids. The LNPs had different compositions, but their apparent pKa as determined by TNS assay gave the same values. In the TNS assay in Figure 2(b and d), the apparent pKa of LP01 and BiP20 LNPs is estimated to be 6.6, making them indistinguishable within our assay. The difference of TNS pKa for LP01 between our results (6.6) and previously reported values (6.1^46^ or 6.3^47^) might be due to some differences between LNP formulations. However, when we studied the interaction of the LNPs with endosomal mimic GUVs, the two LNP formulations showed different pH-dependent interactions.

**Figure 2:**
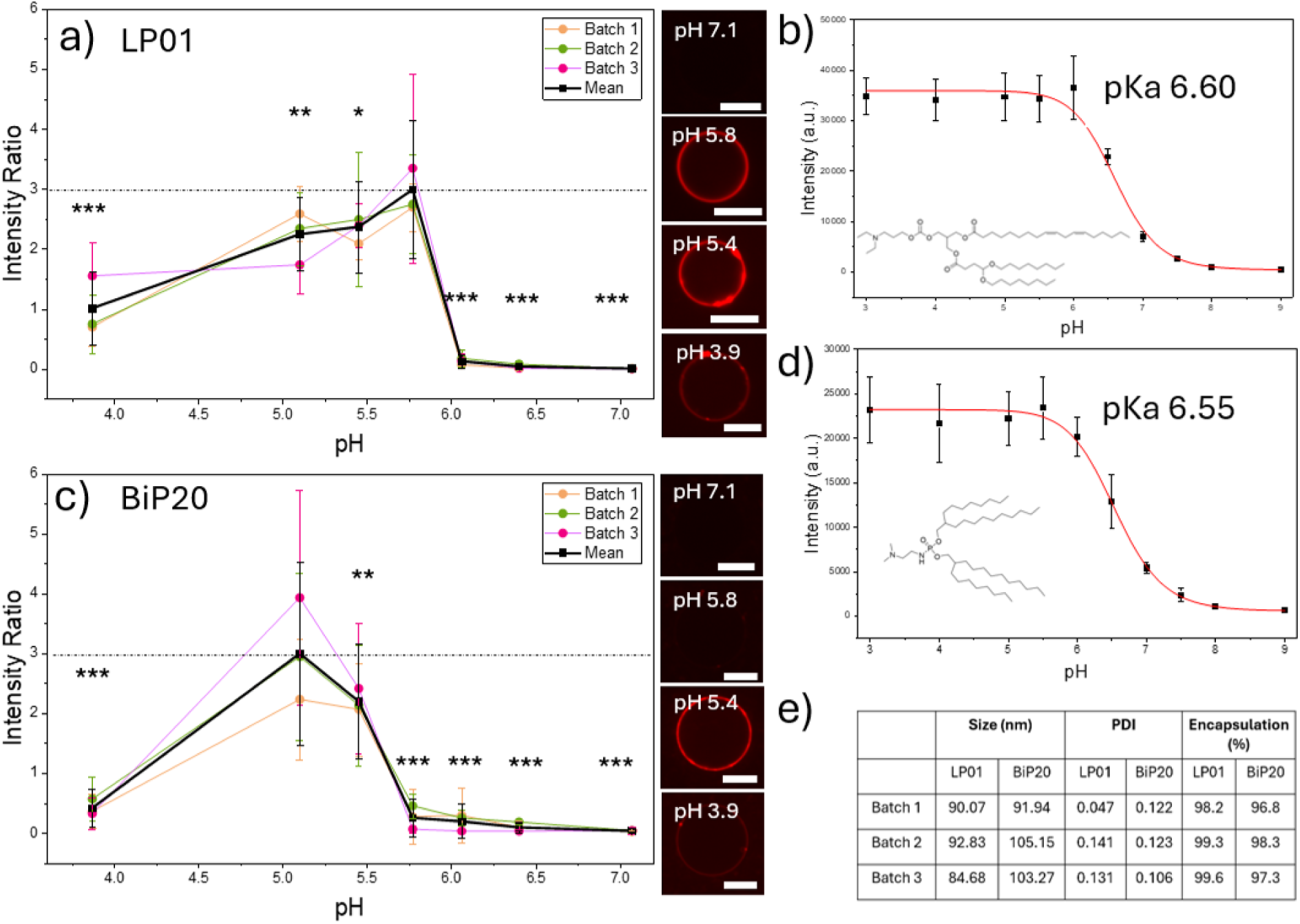
GUVs composed of endosomal lipid mixture are exposed to LNPs made of different ionizable lipids, LP01 and BiP20, at different pH. The LNPs encapsulated Fluc mRNA and were doped with fluorescent lipid Rh-DHPE, and the final mRNA dosage in all samples is 720ng. a) Endosome mimicking GUVs exposed to LP01 LNPs. Representative images on the right; b) pKa estimation of LP01 using the TNS assay along with the chemical structure of LP01; c) Endosome mimicking GUVs exposed to BiP20 LNPs. Representative images on the right; d) pKa estimation of BiP20 using TNS assay along with the chemical structure of BiP20; e) Characterization of all LNPs batches. For a and c, the experiments were repeated for n = 3 independent LNP preparations with all data plotted within the graphs with different colors, and the average of all data is plotted in black. Data points represent means of all the data, and vertical error bars indicate Standard Deviation (SD). Statistical analysis: The horizontal dashed line highlights the baseline mean value of the pH with the highest interaction, which serves as the statistical reference group. Asterisks positioned directly above data points indicate a statistically significant difference compared to the reference group. *** denotes p < 0.0001, ** denotes p < 0.005, and * denotes p < 0.01. For b and d, each point corresponds to the mean and standard deviation of n = 3 replicates. All scale bars: 10 µm

GUVs with the endosomal composition described in the previous section were used as the endosomal mimicking membrane, and the experiments were repeated for n=3 independent LNP preparations, ≥5 GUVs per pH point per experiment. As shown in Figure 2(a and c), the interaction at each pH was quantified by taking the ratio of the fluorescence of the membrane and the background, as mentioned in the previous section. By this measure, LP01 LNP exhibited maximal lipid mixing at pH 5.8, consistent with early-to-mid endosomal pH, whereas BiP20 LNP showed maximum interaction at pH 5.1, which is closer to late endosomal pH. This 0.7 pH-unit divergence in the pH of peak membrane engagement, despite identical measured pKa, indicates that TNS-derived pKa alone does not predict the pH at which a given ionizable lipid produces maximal lipid transfer to a target membrane. The TNS assay works by detecting fluorescence when TNS partitions in the hydrophobic LNPs driven by the electrostatic interaction with the charged surface of the LNPs. The LNP surface gets progressively protonated as the pH is reduced, increasing the fluorescence of TNS and giving an indirect readout of just the surface charge based on the surrounding pH^48–50^; the utility of this assay with regard to the ionizable lipids located in the core of the LNP may be limited^33,50^.

This distinction in molecular interactions of the LNPs with the model membrane could be a result of the lipid architecture of the ionizable lipids. LP01 is an ionizable amino lipid, where a single tertiary amine is linked via an ester bond to two hydrophobic tails, without any phosphate-containing backbone^46,51^. This lipid also features several oxygenated functions (carbonate, esters, acetal; Figure 2) that are all hydrogen bond acceptors. BiP20 belongs to a structurally distinct class of branched ionizable phospholipids, in which a tertiary amine is linked to two branched lipid chains via a phosphoramidate moiety acting as a spacer. The presence of the two branched lipid chains can also be viewed as the presence of 4 lipid tails^23^. BiP-20 contains fewer oxygen atoms than LP01 and only one oxygen atom is prone to accept hydrogen bonds (the P=O motif^52^). The TNS assay reports protonation behavior in a standardized reference environment common to both measurements. However, it does not necessarily capture how each lipid’s local electrostatic and hydration environment shifts once the lipid is in proximity to a compositionally complex, anionic (PS- and PI3P-containing) target membrane such as the endosomal mimicking GUV membrane^48^.^51,52^

The interaction between the bilayer-forming phospholipids and LNPs is governed by hydration repulsion, intramolecular electrostatics, hydrogen-bonding, and the presence of poly(ethylene glycol) lipids may affect the lipid’s interaction with the phospholipid of the target membrane^53–55^. This would alter the effect of the ionizability of amine group in the LNP, requiring a more acidic environment to reach the degree of protonation needed for maximal interaction with the endosomal mimic.

This is consistent with the computational work by Trollmann et al showing that the local environment of the ionizable lipid affects how it partitions in a flat membrane, affecting its apparent pKa^40^. Our data of LNP interacting with endosomal mimicking GUVs at different pH provide direct, membrane level experimental evidence for this divergence from the TNS assay. TNS-based pKa screening may be used as a potential indicator of LNP performance, but it does not predict the complete picture of the interaction with the membrane. Such membrane-based assays, though time-consuming, can provide a better estimate of the maximal activity of novel ionizable lipids, particularly for architectures that depart from the conventional amino-lipid scaffold.

Another interesting result is the lowered membrane fluorescence intensity at pH 3.8 in the case of both LNPs involving LP01 as well as BiP20 (Figure 2). As mentioned above, local disturbance on the membrane recruits more LNPs to interact with the membrane. Once the system reaches very low pH, the LNPs may get more positively charged, and electrostatic repulsion inhibits recruiting more LNPs for the interactions^15^. The PS headgroups of the phospholipids in the GUV also lose their anionic charge in 4acidic conditions, thus reducing the electrostatic attraction between the LNPs and GUVs.^56^

### Lipid Nanoparticles incorporate in the membrane and cause morphological changes

To directly visualize the morphological changes caused by the LNPs on the endosomal mimic membrane, we made GUVs with a fluorescent dye DiO and exposed them to LNPs with a fluorescent cargo, FlucCy5. In the previous section, we saw that both LNP populations interact with the endosomal mimicking GUVs at pH 5.5, so we performed our experiments at this pH.

In the GUV membrane, lipids with headgroups PC and PS have close to zero spontaneous curvatures, making them cylindrical shaped, and lipids with the PE headgroup have negative curvature, making them slightly conical shaped^57^. From Figure 3, we can see that in the absence of LNPs, the GUVs form the lipid tubes on the inside. This indicates a net negative spontaneous curvature resulting from the lipid composition and the buffers on both sides of the membrane^58^. Upon addition of LNPs at pH 5.5, LNPs were attracted to and interacted with the GUV membrane, pulling the membrane tubes to the outer GUV interface. The inserted lipids do not cause a drastic increase in the membrane area. When the GUV membrane increases in area, keeping the volume constant, it results in deflated GUV structures or other complicated membrane morphologies owing to the area excess^59–62^. Instead, we observe that the LNP forms discrete lipid clusters on the GUV surface, visible as dark spots in brightfield imaging.

**Figure 3:**
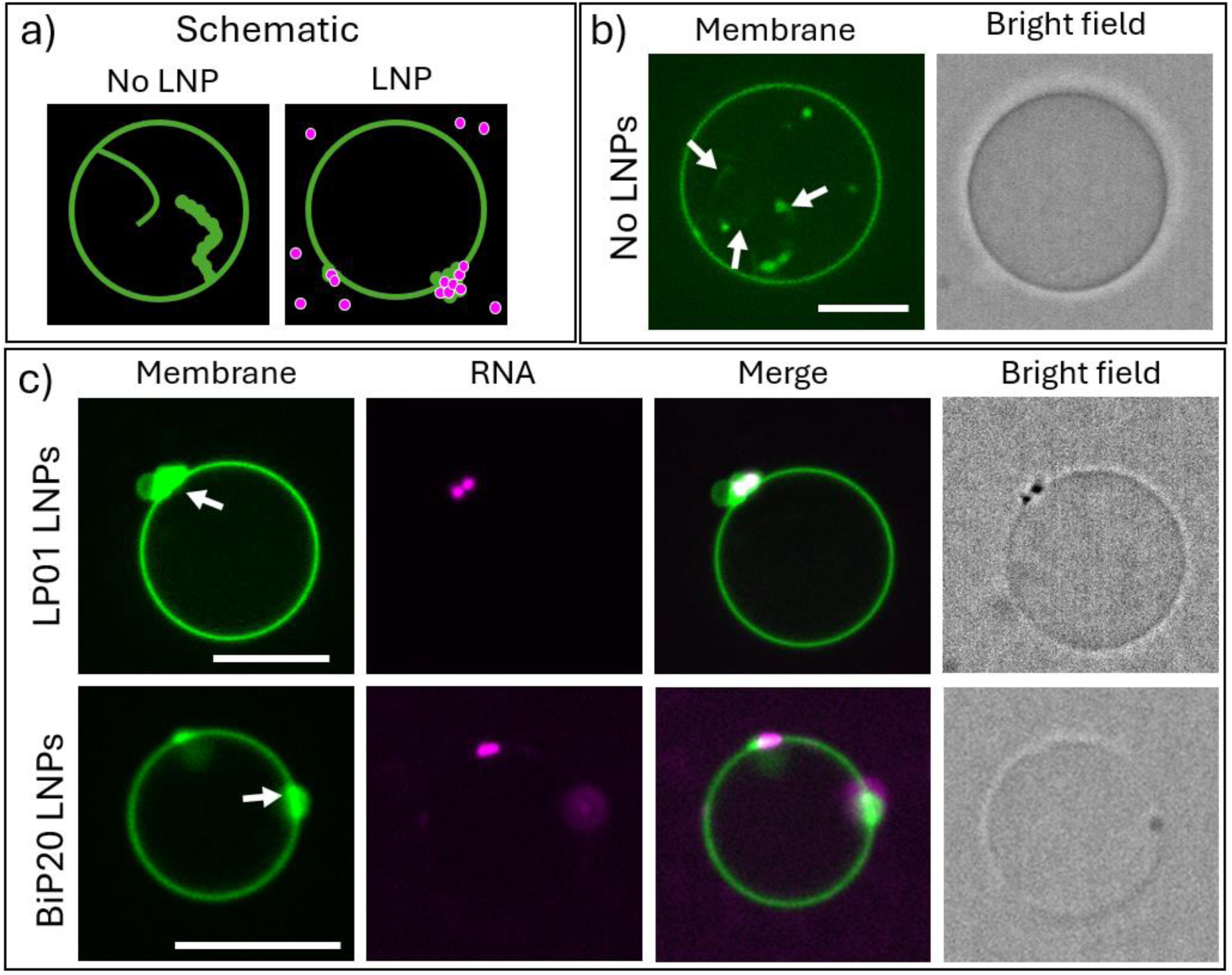
GUVs composed of endosomal lipid mixture, doped with DiO lipid dye (green) and exposed to LNPs made of different ionizable lipids-LP01 and BiP20, encapsulating Fluc-Cy5 mRNA (Magenta). Final mRNA dosage in all samples is 720ng. a) Schematic of the experiment and the results b) In the absence of LNPs, owing to the negative spontaneous curvature, GUVs form tubes on the inside as seen in the fluorescence channel. c) GUVs exposed to LP01 LNPs at pH 5.5. d) GUVs exposed to BiP20 LNPs at pH 5.5 that also form the clump structures on the surface. There are no tubes on the inside, and the membranes form clumps on the GUV surface. These clumps are also visible on the bright field image as dark spots. They coincide with the magenta mRNA signal, showing that the LNPs induce these. All scale bars: 10 µm

Notably, the Cy5-labeled mRNA cargo remained localized to the GUV surface rather than being released into the vesicle lumen. Coarse-grained simulation studies of lipid carrier-membrane fusion show that nucleic acid cargo escape is a slow, stepwise process involving stalk formation, lipid exchange, and water-filled pore formation, rather than immediate bulk cargo release upon contact^63^. Altogether, these results show that the LNPs do transfer some of the helper lipids to the endosomal membrane but do not fuse completely.

### The mRNA is exposed without complete fusion of the membrane

In the previous section, we saw that there is lipid transfer between the LNP and the GUV. To understand what happens to the mRNA cargo, we used a SYBR Gold based imaging assay. SYBR Gold is a highly sensitive fluorescent dye that belongs to the asymmetric cyanine dye family ^64,65^. It is widely used in molecular biology to detect nucleic acids, including double-stranded DNA, single-stranded DNA, and RNA, by intercalating in the nucleic acid backbone. On binding, the dye exhibits a greater than 1000-fold increase in its fluorescence signal^66^. Its high sensitivity, broad nucleic acid compatibility, and favorable photophysical properties have made SYBR Gold a powerful tool for applications such as gel electrophoresis staining, plate reader assays, and fluorescent microscopy^65–67^. To study the exposure and release of mRNA into the surrounding environment, we supplemented the outer solution of the GUVs with SYBR Gold and exposed them to LNPs containing mRNA cargo labeled with Cy5 dye. As seen in Figure 4, the mRNA signal overlaps with the SYBR Gold signal, indicating that the mRNA gets exposed while it is still on the membrane. While the LNP is still on the membrane, the mRNA cargo is exposed, but it does not diffuse away freely. In Figure 4b, we quantified the SYBR Gold intensity as observed on the membrane of the GUVs. At this pH, the mRNA from LP01 LNPs is more exposed to the surroundings, showing more disruption of the LNPs compared to the BiP20 LNPs.

**Figure 4:**
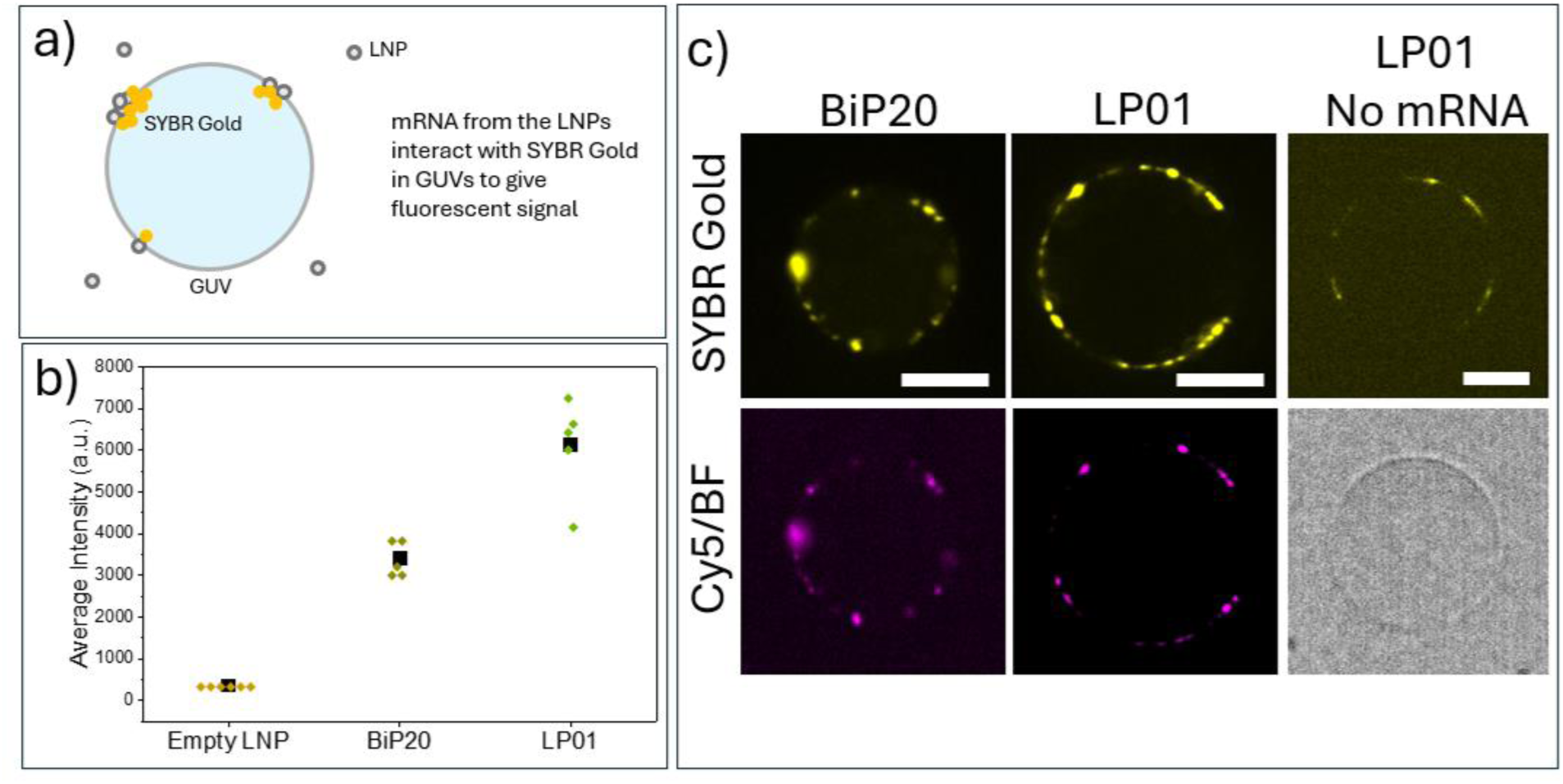
GUVs are exposed to LP01 and BiP20 LNPs encapsulating a mixture of Fluc:Fluc-Cy5 mRNA in the molar ratio 2:3, at pH 5.5 and in the presence of SYBR Gold dye. and without mRNA. a) Schematic of the experiment, where the encapsulated mRNA fluoresces when interacting with SYBR Gold dye. b) Average fluorescence intensities of SYBR Gold localized on the membrane for LNPs with and without mRNA. c) Representative images of mRNA localization on the membrane as seen in the Cy5 channel and the exposure of the mRNA (magenta), SYBR Gold channel (yellow). All scale bars: 10 µm

This is consistent with the computational results that found that the mRNA and the ionizable lipids around it are kinetically bound by electrostatic forces, resulting in the mRNA being stuck near the membrane^16,17^. Kjølbye et al. showed in their simulations that the LNPs induce deformations of the vesicle by forming RNA-membrane clusters, and the helper lipids from the LNP diffuse in the vesicle^16^. Johansson et al. showed colocalization of the ionizable lipids of the LNP with endosomal membrane damage sensor, galectin^11^. We observe mRNA-membrane clusters as well as diffusion of the helper lipids in our experiments shown in Figure 2. During imaging that tool places over 30 mins, we observed membrane junctions in the GUV membrane, similar to ones shown in the study by Kjølbye et al^16^. These vesicles then proceeded to rupture, releasing the mRNA from the lipids, thus mimicking endosomal escape.

Our experiments with GUV and LNPs provide a structural observation that may be relevant to the mechanism of endosomal escape. Figure 5a shows the three membrane pieces joined together, generating regions of high curvatures in the membrane. GUV membranes are in the order of 4-5nm thick and surrounded by aqueous solutions, making it difficult to observe using bright-field microscopy. To confirm that the observed membrane junctions were not artifacts from fluorescent dyes, we labeled the GUVs, LNPs, or mRNA with various dyes in different experiments. We observed similar highly curved hemifusion like morphologies in all experiments. These structures are difficult to capture as the GUVs either burst or move away from the field of view quickly. Despite these experimental limitations, we could capture some of the membrane junctions and a few movies showing the GUV rupture. We believe that these morphologies are a result of the lamellar-to-hexagonal phase transition, resulting from the transfer of the ionizable lipids from the LNPs to the lipid bilayer^68,69^. Examples of these membrane morphologies at different experimental conditions are shown in Figure 5(b-e).

**Figure 5:**
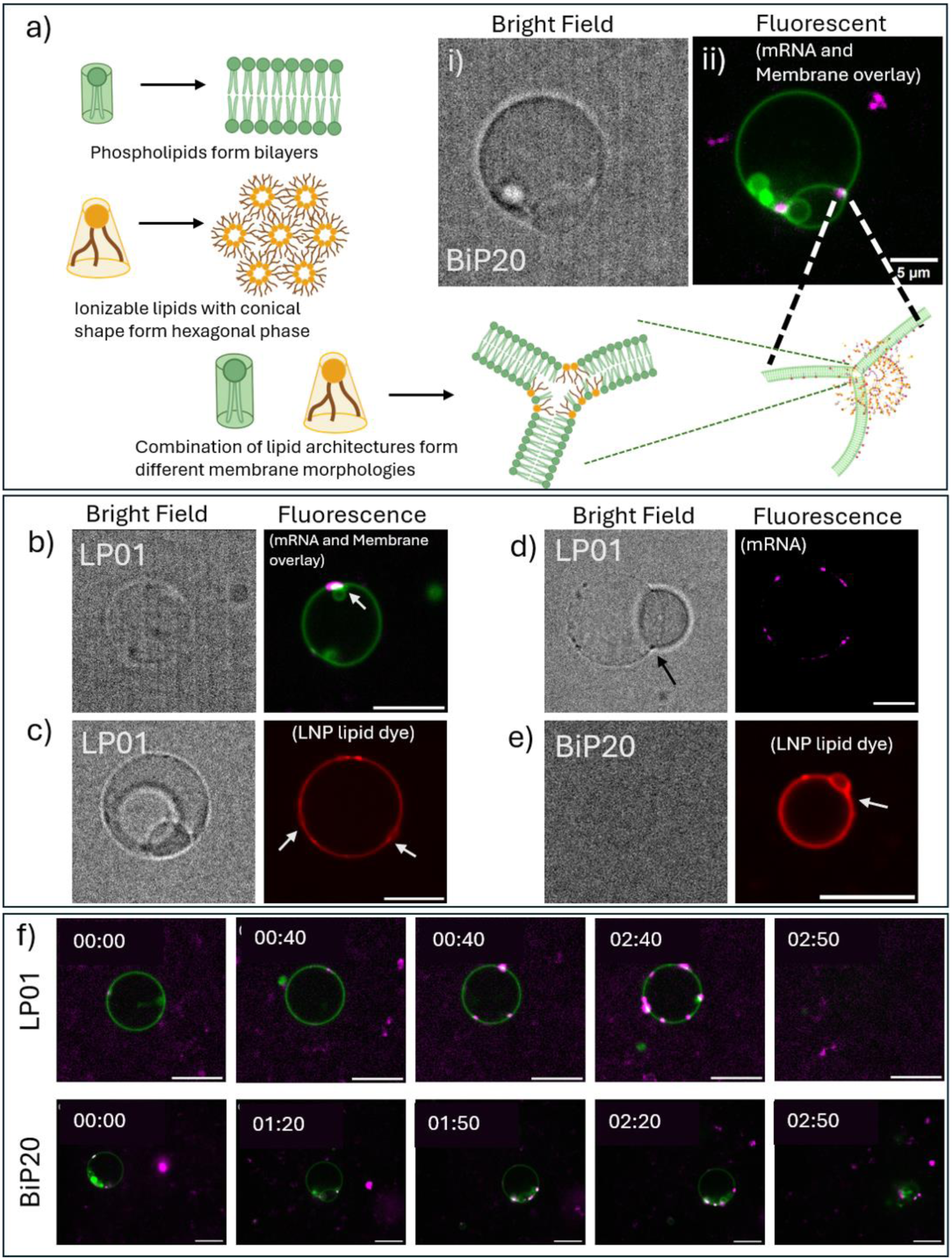
Local membrane disruption caused by LNPs on interacting with the GUVs, resulting in a lamellar-to-hexagonal phase change at the point of interaction. a) Schematic of lipid architecture and their preferred assembly. i-ii) BiP20 LNP encapsulating Fluc-Cy5 (magenta) interacting with GUV with DiO dye (green) at pH 5.5. (b-e) Bright field and fluorescent images of lamellar to hexagonal phase transitions as imaged using different dyes and LNPs made of different ionizable lipids, LP01 and BiP20. b) LNP: LP01, Fluc-Cy5 mRNA (magenta), GUV: DiO membrane dye; c) LNP: LP01, Rh-DHPE lipid dye (red), GUV: no dye; d) LNP: LP01, Fluc-Cy5 mRNA (magenta), GUV: no dye; e) LNP: BiP20, Rh-DHPE (red), GUV: no dye. (f-g) Snapshot from time series recorded following a GUV labeled with DiO membrane dye (green), interacting with LNPs encapsulating Fluc-Cy5 mRNA (magenta). f) LP01 LNP; g) BiP20 LNP. Timestamp format: mm:ss; unless specified, scale bar: 10 µm

This finding suggests that these structures are not artifacts from imaging conditions, but they are a result of the interaction of the endosomal mimicking membrane with the LNPs. The structure of this interface is like hemifusion diaphragms that are observed when two vesicles fuse together^59,70^. Such structures are stabilized by the presence of cone shaped lipids in the highly curved regions^70–73^. Figure 5 shows that the three membranes are joined at these highly curved junctions and are also co-localized by the mRNA signals, indicating the localization of LNPs. The ionizable lipids have broad hydrophobic tails and smaller head groups, making them a favorable lipid to induce highly curved regions. Previous works have clearly shown the propensity of cationic phospholipids, similar to BiP-20 but with a permanent cationic charge, to produce an inverted hexagonal phase^74^. With BiP-20 we hypothesize a similar behavior as soon as it is protonated. This process is consistent with the endosomal escape being mediated by the hexagonal phase transition of the lipids^3,10,75–77^. These structures are sensitive to factors such as the membrane tension, lipid geometries, and the presence of ions or proteins^78–80^. Eventually, the GUVs with LNPs on their surface burst open. Movie snapshots of two such examples of such GUV bursting events caused by LNP interactions are shown in Figure 5(f-g). The complete disruption of the membrane releases the mRNA cargo and completes the process of endosomal escape.

## CONCLUSION

This study used giant unilamellar vesicles mimicking the endosomal membrane (POPC:DOPE:DOPS:cholesterol:PI(3)P) to directly visualize how lipid nanoparticles destabilize the endosomal membrane. This provides a membrane-level mechanistic insight of endosomal escape. We studied two ionizable lipids, LP01 (a conventional amino lipid), and BiP20, a branched ionizable phospholipid (BiP) made in-house. We found that the pKa alone, as measured by standard TNS titration, is insufficient to predict the pH at which the LNP interacts with the membrane. Despite both LNPs sharing an identical measured apparent pKa of 6.6, LP01 showed maximal membrane interaction at pH 5.8, consistent with early endosomal conditions, while BiP20 required the more acidic environment of pH 5.1, characteristic of late endosomes and lysosomes. This divergence, reproducible across independent LNP preparations and multiple GUVs per condition, indicates that molecular LNP properties such as ionizable lipid architecture and helper lipids modulate the functional pKa realized at the membrane interface, independent of the pKa measured using the field-standard TNS assay.

We acknowledge several caveats inherent to this reductionist system. Our GUVs are protein-free and therefore do not recapitulate the biomolecular protein corona that LNPs acquire in vivo; although this corona may partially dissociate during cellular uptake and endosomal trafficking, its presence could still modulate the LNP-membrane interactions reported here, and LNP size and morphology may likewise differ under physiological conditions from the formulations characterized in vitro. In addition, LNPs in our experiments engage the outer leaflet of the GUV, modeling interaction with the outer limiting membrane rather than the luminal inner leaflet that an LNP would encounter within an intact endosome. We recognize these limitations and view the defined GUV platform as a complement to, rather than a replacement for, other cellular and in vivo studies.

At the membrane level, LNP engagement produced a sequence of morphological changes. The membrane tubes formed at the inner interface of GUVs were first pulled out, with the LNPs clustering in specific regions in the membrane instead of homogenous interaction with the membrane. This led to the formation of hemifusion-diaphragm-like junctions followed by the lysis of the membrane. Notably, across these morphological changes, Cy5-labeled mRNA cargo remained close to the GUV membrane surface rather than being released into the vesicle lumen. This observation suggests that membrane destabilization and cargo release are mechanistically separate steps. The diaphragm stage acts as an intermediate step that incorporates the ionizable lipids in the endosomal membrane, triggering a hexagonal lipid phase. This transition itself does not release the mRNA cargo. Our images and data point towards a complete lysis of the endosomal membrane leading to the release of mRNA in the cytosol. Such GUV-based studies offer a cell-free platform for isolating each of the mechanistic steps of interactions of LNPs with endosomal membranes. This is particularly valuable for structure prediction and evaluation of next-generation ionizable lipid chemistries whose functional behavior can diverge substantially from what bulk biophysical characterization predicts.

## MATERIALS AND METHODS

### GUV preparation

GUVs were made using PVA gel swelling method. 4% PVA solution was prepared by dissolving 40 mg of PVA in 1 mL of water. 15 μL of this solution was spread on clean cover glass (1.5cm x 1.5cm) and dried in an oven at 90 °C for 10 min. Using a glass syringe, 3 μL of 1.5 mM lipid mixture in chloroform was smeared on the dried PVA and the chloroform allowed to evaporate at room temperature for 10 min. The cover glass was placed inside a well of a 6-well plate. With a pipette, 900µL of 300 mM sucrose in 2 mM HEPES buffer was gently placed over the cover glass, so that nothing spilled over the coverglass. The setup was covered and left undisturbed for an hour at room temperature. The vesicles were harvested by gentle pipetting and stored at room temperature till further experiment. Vesicles were prepared fresh before every experiment.

All lipids were purchased dissolved in chloroform from Avanti Lipids, with the exception of cholesterol, which was purchased from Sigma-Aldrich.

### LNP preparation

mRNA-encapsulated LNPs were formulated by microfluidic mixing of lipid and mRNA phases using a NanoAssemblr Benchtop Instrument (Precision Nanosystems). The lipid phase consisted of ionizable lipid, phospholipid, cholesterol, and PEG-lipid dissolved in ethanol at one of two formulations: LP01/DSPC/cholesterol/DMG-PEG2k (45.5:9:44:1.5 mol%) or BiP20/DOPE/cholesterol/DMPE-PEG2k (50:9.5:38.5:1.5 mol%). In some experiments, the helper lipid in both formulations was substituted with the fluorescent lipid Rhodamine-DHPE at 0.5 mol% of the total lipid composition.

All lipids were purchased from Avanti Lipids, with the exception of cholesterol, which was purchased from Sigma-Aldrich, DMG-PEG2k that was purchased from NOF America Corporation. BiP-20 was prepared as previously reported by using as a key reaction the Atherton-Todd reaction to react bis(2-octyldodecyl)phosphite with *N*,*N*-dimethylethylenediamine^23^.

The aqueous phase consisted of mRNA (Fluc, Fluc-Cy5 or a mixture of Fluc:Flu-Cy5 in the molar ratio 2:3) in 100 mM citrate buffer (pH 4.0). Lipid and aqueous phases were combined at a 1:3 flow rate ratio (organic:aqueous) per the manufacturer’s protocol. Following formulation, LNPs were diluted in 1× DPBS (pH 7.4) and dialyzed against 3 L PBS using 10 kDa MWCO Slide-A-Lyzer cassettes (Thermo Fisher Scientific) for 3 h at room temperature, then transferred to fresh PBS and dialyzed overnight at 4°C. Dialyzed LNPs were concentrated using 10 kDa MWCO Amicon Ultra-4 mL centrifugal filter units (Millipore, Burlington, MA).

### LNP characterization

Particle size and polydispersity index (PDI) were measured by Stunner (Unchained Labs) at 25°C. Encapsulation efficiency and mRNA concentration were quantified using the Quant-iT RiboGreen RNA Assay Kit (Thermo Fisher Scientific, Waltham, MA).

### TNS Assay

LNPs were diluted to 40 μg ml^−1^ of encapsulated nucleic acid in 1× PBS. In a black 96-well plate, 90 μl of unique pH buffers, ranging from pH 3.50 to pH 9.0 and 3.26 μl of dilute LNP solution was added. Finally, 2 μl of 300 μM TNS reagent was added to each well. All data points were prepared in triplicates. The plate was set on a plate rocker for 5 min, from which the fluorescence values were read (excitation 325 nm, emission 435 nm). The fluorescence emission data was fitted to sigmoid, and the point of inflection was noted to be the apparent p*K*_a_ of the LNP formulation.

### Image acquisition and analysis

All fluorescent images were recorded using a spinning disc confocal microscope (Nikon). Excitation wavelength of 532 nm was used to excite Rhodamine, 477 nm to excite SYBR Gold and 637nm to excite Cy5. The emission was collected using appropriate filter cubes. All images were taken using a 60x oil immersion objective.

The radial profile angle plugin of Fiji was used to quantify the intensity on the membrane. To avoid reading from membrane clusters, instead of the entire vesicles, the radial profile was plotted for regions on the vesicles without clusters for an angle between 60° and 120°, as shown in Figure 1. The membrane intensity was calculated by first subtracting the background using ‘peak analyzer’ in OriginPro 2025b and then integrating over the whole radial range. The background intensity was averaged over three regions in the background, avoiding membrane clusters. Finally, the ratio of the membrane intensity and background intensity was taken. For SYBR Gold intensity measurement, threshold brightness of 2000 was applied to all images of LNPs with mRNA to select the regions of interest. The average intensities of them were noted for each of the GUV.

Statistical comparisons across pH conditions (Figure 2) were performed using one-way ANOVA followed by Tukey’s post-hoc test for pairwise multiple comparisons, using OriginPro 2025b. Asterisks indicate statistical significance relative to the reference group (the pH condition with the highest mean interaction): *** p < 0.0001, ** p < 0.005, * p < 0.01.

Fluc was purchased from Helix, and Fluc-Cy5 was made in-house using the protocol mentioned below.

### Synthesis of Fluc-Cy5 mRNA

Labeled mRNA Fluc-Cy5 was synthesized in-house following these steps.

#### DNA Template Synthesis

The untranslated regions used for mRNA construction were derived from our previous work^81^. A codon-optimized DNA sequence encoding Photinus pyralis firefly luciferase (Fluc; GenBank accession AAA29795.1) was synthesized by GenScript. Linear DNA templates for in vitro transcription were assembled by fusion PCR using Q5 Hot Start High-Fidelity DNA Polymerase (New England Biolabs). Each template contained, from 5′ to 3′, a T7 promoter, the 5′ untranslated region, the Fluc open reading frame, the 3′ untranslated region, and a 120-nt poly(A) tail encoded in the reverse primer. PCR products were purified using the DNA Clean & Concentrator-5 kit (Zymo Research), verified by agarose gel electrophoresis, and quantified using a NanoDrop spectrophotometer (Thermo Fisher Scientific).

#### Synthesis and Purification of mRNA

Fluc mRNA was synthesized from the linear DNA template using the HiScribe T7 High Yield RNA Synthesis Kit (New England Biolabs), with CleanCap Reagent AG (3′ OMe) (TriLink BioTechnologies) included for co-transcriptional capping, according to the manufacturers’ instructions. After transcription, the residual DNA template was removed by treatment with TURBO DNase (Invitrogen). The RNA was purified using the Monarch RNA Cleanup Kit (New England Biolabs) and eluted in nuclease-free water. The mRNA concentration was determined by measuring absorbance at 260 nm using a NanoDrop spectrophotometer (Thermo Fisher Scientific). Transcript size and integrity were assessed by denaturing urea-polyacrylamide gel electrophoresis. Purified mRNA was aliquoted and stored at −80 °C until use.

#### Cy5 Conjugation to mRNA

Cy5 was conjugated to the 3′ end of the mRNA using pCp-Cy5 (1 mM stock solution; Jena Bioscience) and T4 RNA Ligase 1, High Concentration (New England Biolabs). Each labeling reaction contained 150 μg of mRNA in a total reaction volume of 600 μL, with 1× T4 RNA Ligase Reaction Buffer, 1 mM ATP, 10% DMSO, 10 μM pCp-Cy5, 900 U of T4 RNA Ligase 1, High Concentration, and 240 U of RNase Inhibitor, Murine (New England Biolabs). The reaction was incubated at 16 °C for 16 h protected from light. Cy5-conjugated mRNA was subsequently purified using the Monarch RNA Cleanup Kit (New England Biolabs) and stored at −80 °C until use.

## AUTHOR INFORMATION

### Author Contributions

SP: Conceptualization, Methodology, Data curation, Formal analysis, Writing-Original Draft and Editing; AV and AK: LNP Formulation and Characterization; ZL: mRNA-Cy5 Synthesis; MB and P-AJ: Synthesis of BIP20 lipids, Writing-Review and Editing; GS and YE: Conceived of the idea and directed research, Resources, Funding acquisition, Writing-Review and Editing; All authors have read the manuscript and approved the final version of the manuscript.

